# High CO_2_ levels overcome the lethality of circadian mutants in cyanobacteria

**DOI:** 10.1101/2025.06.01.657271

**Authors:** Alfonso Mendaña, María Santos-Merino, Marina Domínguez-Quintero, Raquel Gutiérrez-Lanza, Daniel C. Ducat, Raúl Fernández-López

## Abstract

Cyanobacteria have adapted to daily fluctuations in light intensity through a circadian clock that aligns their physiology and metabolism to the external daytime. In the model cyanobacterium *Synechococcus elongatus* PCC 7942, the rhythm generated by a molecular pacemaker is converted into a global transcriptional oscillation by the central regulator RpaA. Mutants in RpaA and other components have been instrumental in elucidating the clock’s molecular architecture and physiological role. Extensive evidence suggests that one of the central functions of circadian regulation is to prevent the accumulation of reactive oxygen species (ROS) during nighttime. However, circadian mutants are unable to grow under natural day/night cycles and require constant illumination, hindering our ability to study the clock function during the night, a phase where circadian regulation is critical for redox homeostasis. Here, we show that the darkness lethality phenotype of circadian mutants can be overcome by high CO_2_ levels. When grown under a 3% CO_2_ atmosphere, RpaA-null mutants exhibited growth rates similar to the wt. An analysis of the ROS levels under different CO_2_ and light intensity conditions revealed carbon scarcity to be the most significant contributor to redox stress. Nighttime ROS accumulation can be modulated by CO_2_ abundance, an observation that will allow the characterization of heretofore lethal mutants in circadian regulation. The dispensability of the circadian clock in a high CO_2_ environment suggests that the clock may have evolved as an adaptation to the decrease in atmospheric CO_2_ levels that occurred after the Great Oxygenation Events, in the Paleoproterozoic era.

**IMPORTANCE:** Cyanobacterial circadian clocks are considered essential for survival under natural day/night conditions, and many circadian mutants are lethal under diel cycles. Results presented here challenge this assumption, showing that these mutants grow normally when CO_2_ is abundant. This suggests that the major adaptive role of the clock is to manage the redox stress caused by carbon limitation. By separating the effects of carbon availability from circadian control, we identify a key environmental factor that shaped the evolution of biological timekeeping. Using high CO_2_ to bypass clock dependency will allow the study of circadian function in otherwise lethal mutants. These findings reframe the clock as a specific adaptation to atmospheric carbon, stressing its key role in the regulation of the redox balance.

## MAIN

Circadian clocks coordinate the physiology of eukaryotic and prokaryotic organisms with the daily fluctuations in light intensity caused by earth’s rotation. The cyanobacterium *Synechococcus elongatus* PCC 7942 (hereafter *Synechococcus*) is a model organism for the study of circadian control because it contains one of the simplest circadian clocks known to date. The cyanobacterial clock consists of a post-transcriptional oscillator formed by the proteins KaiABC, which are able to sustain a 24-hour phosphorylation cycle (1). The clock synchronizes its internal state with external time through two light-dependent cues: the ATP/ADP ratio, and the redox state of the photosynthetic electron transfer chain (2). The timing of the clock is then converted into transcriptional rhythms by the SasA-RpaA two component system, which regulate most genes of the genome (3–5). The adaptive role of the cyanobacterial circadian clock under physiological light/dark (LD) conditions has been established by a combination of genetic, physiological and ecologic studies (6). Mutants lacking key components such as the SasA histidine kinase (3) or the output regulator RpaA (4) have been shown to suffer from severe growth deficits or complete non-viability in LD conditions. In LD conditions, wild-type strains outperform not only arrhythmic mutants, but also rhythmic variants whose endogenous free running periods are misaligned with the external 24h rhythm (6). These findings highlight the adaptive role of the clock in anticipating environmental changes and suggest that fluctuations in light intensity represent the key parameter to which the clock responds to.

However, while light is the most obvious environmental cue in diel conditions, this does not necessarily imply that the changes in light intensity represent the environmental challenge to which the clock is adapting to. There is growing evidence that the metabolic consequences of light/dark transitions pose the most significant selective pressure. In *Synechococcus*, mutants lacking RpaA or deficient in SasA function are able to grow under continuous illumination, but exhibit a darkness-induced lethality phenotype, by which exposure to a period of darkness at any point in their cycle results in a sharp loss of cell viability (3, 4). In RpaA-null mutants, the onset of darkness induces the accumulation of lethal ROS levels, as these mutants are unable to mobilize their glycogen reserves to build up NADPH levels to counteract ROS accumulation (7). Oxidative stress is thus the direct cause of the lethality of RpaA-null strains, and a likely cause of the fitness deficit exhibited by mutants with misaligned circadian control.

In plants, atmospheric carbon limitation is a major source of photorespiration and oxidative stress (8), thus we wondered whether carbon availability would have an impact on the darkness-induced lethality of circadian mutants. To test this end, we tested the response to carbon in the wt and two mutants that show impaired growth in LD, a *ΔrpaA* strain and C11. C11 is a strain that was obtained after long-term evolution under continuous illumination, which resulted in a dramatic increase in the growth rate under constant light, but also caused impaired growth when cells are exposed to diel cycles. This phenotype was linked to a mutation in SasA causing an insufficiency in active RpaA levels (9). When the wt, *ΔrpaA* and C11 were grown on BG11 agar plates under an atmospheric CO_2_ concentration (0.04%), 30 °C, and 180 µmol photons m^-2^ s^-1^, we observed a decrease in viability both in *ΔrpaA* and C11 (Figure 1A), a defect that has been previously shown for RpaA null mutants (4, 10). When the same strains were grown in liquid BG11, at 0.04% CO_2_, 30 °C and 120 µmol photons m^-2^ s^-1^, we observed a severe growth deficit in *ΔrpaA* (Figure 1B, purple line) and a slight growth deficit in C11 (Figure 1B, yellow line). Increasing the light intensity to 983 µmol photons m^-2^ s^-1^ resulted in a much more severe growth arrest that now extended even to the wt (Figure 1C). When we repeated the same experiment bubbling the cultures with a 3% CO_2_ atmosphere, we observed that all strains displayed vigorous growth, both at 120 µmol photons m^-2^ s^-1^ and 983 µmol photons m^-2^ s^-1^. Under these conditions, the *ΔrpaA* grew at a faster pace than the wt.

**Figure 1.**
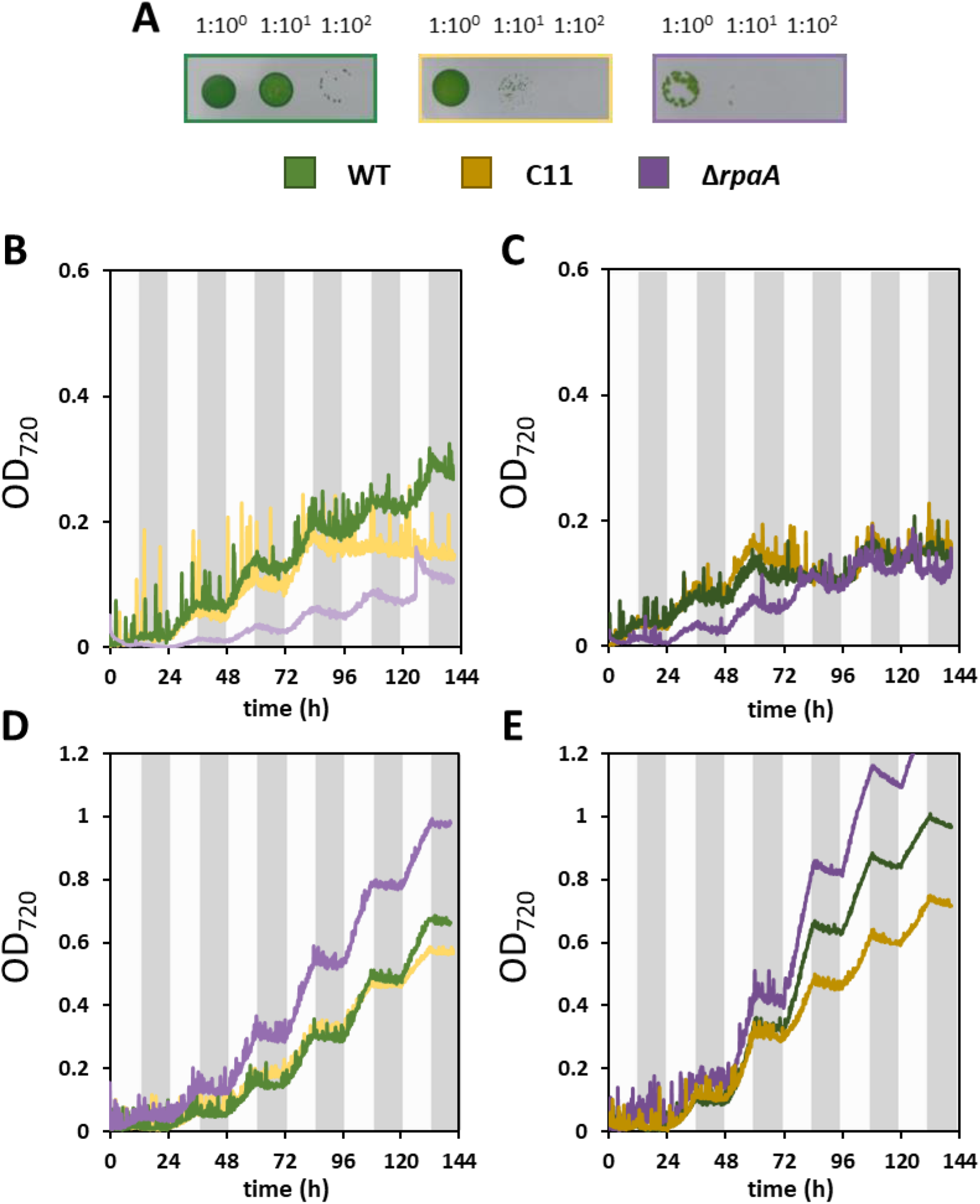
Circadian mutants grow in diel conditions under high CO_2_ atmospheres. A) Growth on BG11 agar plates of serial dilutions of the WT (green), C11 (yellow) and ΔrpaA (purple) under 12h/12h Light-Dark (LD) conditions and 0.04% CO_2_ concentration. B-E) Growth curves obtained under LD on liquid BG11 by the wt (green), C11(yellow) and ΔrpaA (purple). Growth was monitored as OD_720_ along time. Cultures were grown under 0.04% CO_2_ concentrations (panels D, E) or 3% CO_2_ (panels B, C) and a regime of 12 h light and 12 h darkness (grey areas). Light intensity was set at 120 µmol photons m^-2^ s^-1^ (panels B, D) or 983 µmol photons m^-2^ s^-1^ (panels C, E).

Since ROS accumulation has been shown to be the key mechanism behind the darkness-induced lethality observed in *ΔrpaA*, we wondered whether increased CO_2_ concentrations may buffer the redox stress suffered by circadian mutants. For these reasons, we quantified the amount of ROS present in cultures that have been grown under atmospheric (0.04%) or high (3%) CO_2_ concentrations. We first measured the ROS production in cultures grown in BG11, at 30 °C, under continuous illumination at 120 µmol photons m^-2^ s^-1^ and 983 µmol photons m^-2^ s^-1^. ROS levels were measured using the protocol by Velikova et al. (11), as described in Materials and Methods. As shown in Figure 2A, under atmospheric (0.04% CO_2_) conditions, all strains showed similar ROS levels, which significantly increased when cells were shifted from 120 to 983 µmol photons m^-2^ s^-1^. When cells were grown in high CO_2_, under these light intensities, we observed a sharp decrease in ROS levels, which became nearly undetectable in all strains (Figure 2A). Increasing light intensity from 120 to 983 µmol photons m^-2^ s^-1^ caused a slight but significant increase in ROS in the WT and C11, whereas this increase was more pronounced in *ΔrpaA*. Overall results demonstrated that CO_2_ availability is a major driver of ROS accumulation in *Synechococcus*, while light intensity plays a secondary role.

**Figure 2.**
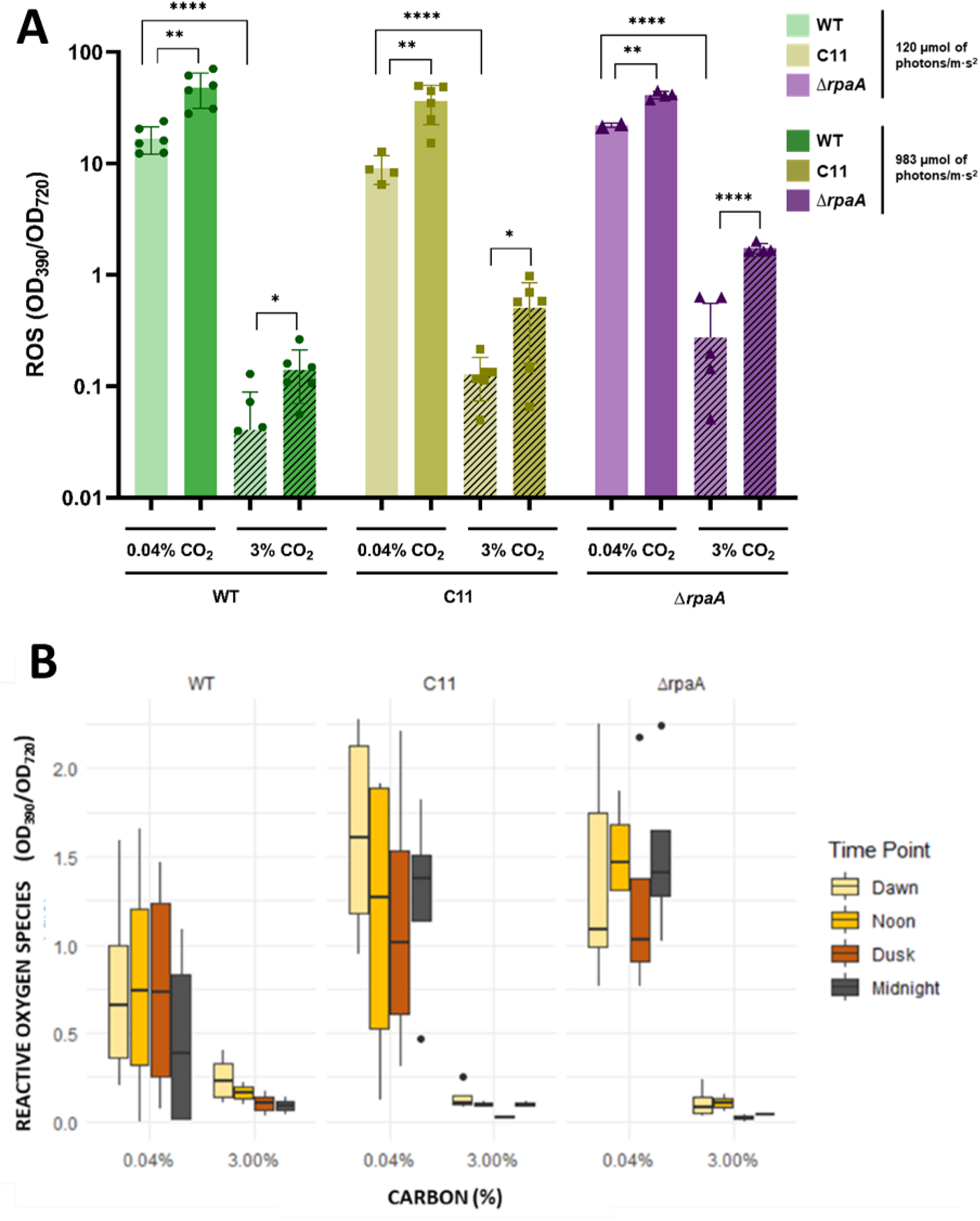
Reactive oxygen species accumulation differs under atmospheric or high CO_2_ conditions. A) Reactive oxygen species determined in the wt (green), C11(yellow) and ΔrpaA (purple) after 48 h of growth under continuous light, in the conditions indicated in the legend . Boxes represent the average of at least 4 independent biological replicates. Statistical significance was calculated using unpaired Student’s t-test. *p-value < 0.05, **p-value < 0.01, ****p-value < 0.0001. B) Reactive oxygen species obtained after five days of growth in diel conditions (12 h L/12 h D) in the wt (right), c11 (middle) and ΔrpaA (right panel). Each box shows the ROS measured at 6 h intervals (dawn, noon, dusk and midnight) over a time course of 24 h. Leftmost panels correspond to cells growth under 60 µmol photons m^-2^ s^-1^ light intensity and 0.04% CO_2_. Right panels for all mutants correspond to ROS levels obtained at 60 µmol photons m^-2^ s^-1^ light intensity and 3% CO_2_

Previous experiments have shown that the increase in ROS levels in *ΔrpaA* mutants occurs specifically during the night, after the onset of darkness (7). For this reason, we measured ROS levels at different time points during a diel cycle in the wt, and the *ΔrpaA* and C11 mutants. Since these mutants show decreased viability in LD cycles at light intensities above 120 µmol photons m^-2^ s^-1^, we opted to perform the experiment at a light intensity of 60 µmol photons m^-2^ s^-1^, which was previously shown to allow the growth of *ΔrpaA*. Cells were grown at this light intensity, under atmospheric (0.04%) CO_2_ concentrations, 30 °C and a LD period of 12h L/ 12hD. ROS levels were extracted at times 0 h (dawn), 6 h (noon), 12 h (dusk), 18 h (midnight) and 24 h (dawn) as described in materials and methods. As shown in Figure 2B, under these conditions we detected a 2-fold mean increase in ROS levels in *ΔrpaA* and a 1.9-fold mean increase in C11 with respect to the wt. Interestingly, results also showed that while ROS levels go down during the night in the wt, they persist in *ΔrpaA* and C11, supporting the idea that nighttime ROS clearance is impaired in circadian mutants (7). When the same experiment was performed under 3% CO_2_, a sharp decrease in ROS was observed at all time points for all strains, a decrease that was maximal in the *ΔrpaA* strain (18.75-fold). Results thus demonstrated that high CO_2_ levels prevent the accumulation of ROS under natural diel cycles, both in the wt and in the circadian mutants.

Altogether, results indicate that high CO_2._ concentrations prevent the accumulation of ROS during light/darkness transitions, and in such conditions the circadian cycle is no longer essential in cyanobacteria. Although these CO_2._ levels are unrealistic for modern times; cyanobacteria are likely to have experienced such concentrations during their early evolution. Different estimates indicate that, in the early stages of photosynthesis evolution during the Archean and Paleoproterozoic eons, atmospheric CO_2_ concentrations were 10 to 250-fold higher than modern levels (12). After the first Great Oxygenation Event, atmospheric CO_2_ steadily declined for the next 1500 million years, reaching its minimum during the Carboniferous, approximately 350 million years ago. As cyanobacteria had to adapt to declining CO_2_ and increasing O_2_ levels, the earth rotation speed slowed down, increasing the length of the day from 10-14 hours to its modern 24 h period. It is thus likely that cyanobacteria experienced increasing oxidative shifts during light/dark transitions, and adapting to such redox stress periods may be the major adaptive advantage of the circadian clock.

## CONTRIBUTIONS

Alfonso Mendaña, Conceptualization, Data Curation, Formal analysis, Investigation, Methodology, Validation, Visualization, Writing – Original draft, Writing – Review & editing | María Santos-Merino, Investigation, Methodology, Validation, Resources, Writing – Review & editing | Marina Domínguez-Quintero, Investigation, Validation| Raquel Gutiérrez-Lanza, Investigation, Validation | Daniel C. Ducat, Funding acquisition, Resources,| Raúl FernándezLópez, Funding acquisition, Project administration, Resources, Supervision, Visualization, Writing – Original draft, Writing – Review & editing.

## MATERIALS AND METHODS Cyanobacterial strains and culture growth conditions

*Synechococcus elongat*us PCC 7942 (PCC 7942) and all derived strains were routinely grown and maintained in liquid BG11 medium supplemented with appropriate antibiotics. Antibiotics used for selecting PCC 7942 were neomycin at 5 or 25 µg/ml (Neo5 or Neo25) and streptomycin at 10 or 50 µg/ml (Sm10 or Sm50). Except otherwise stated, cells were grown under a continuous light flux of 60 µmol photons m^-2^ s^-1^ from white fluorescent lamps, in a Sanyo Plant Growth Chamber, at 30 °C and constant ambient air supply. Experiments with different light intensities, diel cycles and CO_2_ concentrations in liquid cultures were performed in a MC 1000-OD Multicultivator (Photon Systems Instruments), equipped with programmable temperature, CO_2_ and light intensity (cold white LED).

Experiments of diel cycles growth on solid plates were performed in an Ibercex chamber under 12 h of 180 µmol photons m^-2^ s^-1^ illumination followed by 12 h of darkness and constant ambient air supply. BG11 solid plates were obtained from a 1:1 mixture of liquid 2x BG11 medium and a 2% w/v bactoagar mix and were supplemented with 0.05 mM HEPES and the appropriate antibiotics. Cells were grown under continuous illumination until reaching OD_720_ = 0.6 and then, were serially diluted and 10 µl of each dilution were plated three times per plate.

### Genetic constructions

Genetic engineering in PCC 7942 was performed through natural transformation. For each transformation, approximately 4×10^9^ cells, equivalent to 10 μg of chlorophyll, were mixed with 500 ng of DNA, and incubated in the dark for 16h at 30 °C and 100 rpm. Transformation mixtures were then deposited onto a 0.45 μm nitrocellulose filter (Millipore) and incubated for 24 h on top of BG11 plates supplemented with appropriate antibiotics, at 30 °C and continuous light. The filter was transferred to new plates with antibiotics every 24-72 hours. Transformation colonies appeared after 7-14 days. Segregation of mutants was achieved by repeatedly striking individual transformant colonies on selective plates. Mutant genotypes were confirmed by PCR and DNA sequencing using specific primers.

### Reactive oxygen species colorimetric determination

To measure the total amount of ROS produced by the different strains we used as proxy the yellow product formed in the reaction between potassium iodide (KI) and H_2_O_2_, produced by the cell, adapted from the method by Velikova et al (11). 100 μl of 100% trichloroacetic acid were mixed with 900 μl of culture and centrifuged at 17,000xg 11 min. 500 μl of the supernatant were mixed with 250 μl of phosphate buffer 10 mM and 250 μl of KI 1 M and inverted 3-6 times to properly mix the solutions. After a 20-25 min incubation in the darkness, Abs_390_ was measured and divided by the OD_720_ to normalize the data.

